# An iterative and automated computational pipeline for untargeted strain-level identification using MS/MS spectra from pathogenic samples

**DOI:** 10.1101/812313

**Authors:** Mathias Kuhring, Joerg Doellinger, Andreas Nitsche, Thilo Muth, Bernhard Y. Renard

## Abstract

Untargeted accurate strain-level classification of a priori unidentified organisms using tandem mass spectrometry is a challenging task. Reference databases often lack taxonomic depth, limiting peptide assignments to the species level. However, the extension with detailed strain information increases runtime and decreases statistical power. In addition, larger databases contain a higher number of similar proteomes.

We present TaxIt, an iterative workflow to address the increasing search space required for MS/MS-based strain-level classification of samples with unknown taxonomic origin. TaxIt first applies reference sequence data for initial identification of species candidates, followed by automated acquisition of relevant strain sequences for low level classification. Furthermore, proteome similarities resulting in ambiguous taxonomic assignments are addressed with an abundance weighting strategy to improve candidate confidence.

We apply our iterative workflow on several samples of bacterial and viral origin. In comparison to non-iterative approaches using unique peptides or advanced abundance correction, TaxIt identifies microbial strains correctly in all examples presented (with one tie), thereby demonstrating the potential for untargeted and deeper taxonomic classification. TaxIt makes extensive use of public, unrestricted and continuously growing sequence resources such as the NCBI databases and is available under open-source license at https://gitlab.com/rki_bioinformatics.

## Introduction

Pathogenic strain identification using LC-MS/MS-based proteomics presents a crucial, yet highly challenging task. Many pathogenic strains feature significant phenotypic differences within a species with respect to pathogenicity, zoonotic potential, cell attachment and entry, host-virus interaction and clinical symptoms ^1–3^. In a diagnostic context, strain-level knowledge is important to infer virulence ^4,5^ and drug resistance ^6^ for appropriate therapy. However, inferring exact strain information from proteomic samples remains a difficult task, in particular when the taxonomic origin of a sample is unknown and when related strains feature high sequence similarity.

In recent years, MALDI-TOF mass spectrometry has gained popularity as fast, sensitive and economical method for microbial biotyping. However, identifying strains using MALDI-TOF workflows is still very challenging and requires curated, often proprietary spectral databases ^7^. Several commercial platforms for microbial biotyping down to the species or strain level are available based on MALDI and other technologies such as the Bruker MALDI Biotyper Systems ^8^, the Bruker Strain typing with IR Biotyper ^9^ and the Ibis T5000 Universal Biosensor ^10^.

Several studies report on limitations of MALDI-TOF biotyping for strain-level identifications and advocate advancements towards MS/MS marker peptide detection and, consequently, the analysis was shifted to the MS/MS level ^11,12^. In these studies, however, the MS/MS-based protein identification was established using sequence databases that were already targeted or restricted to particular species or limited sets. In contrast, untargeted MS/MS typing approaches are limited to species level identification ^6,13^. However, in general MS/MS is preferred for the analysis of complex unpurified peptide mixtures as it is considered to provide more distinct and unambiguous peptide and protein identifications ^14^ and thus increased proteome resolution ^15^ as well as higher statistical confidence ^16^. In particular, organisms with unknown taxonomic origin benefit from peptide sequence-based analysis, as MALDI-based biotyping is in comparison too prone to ambiguous identifications ^17^. Furthermore, advances in instrumentation including higher resolution, mass accuracy and dynamic range increasingly allow for identification of the majority of all fragmented peptides ^18^ resulting in higher sensitivity, higher coverage of target proteomes and thus higher availability of distinctive features.

Taking advantage of the vast amount of available protein sequences for MS/MS strain-level identification is challenging. On the one hand, constraining the search space may result in unidentified strains or incorrectly assigned taxa, in particular for non-model organisms ^19^. On the other hand, applying large databases is not recommended either since it decreases peptide identification rates ^20^ and thus eventually impedes taxonomic inference ^21^. Furthermore, with increasing database size sequence quality often decreases (e.g. when using the complete NCBI Protein in comparison to the NCBI RefSeq database) and contaminant sequences may occur more often ^22^. Therefore, extended databases should only be used when necessary. However, strain-level identification of MS/MS spectra from samples with unclear taxonomic status requires an untargeted search against comprehensive databases holding as many strains as possible.

A common and popular concept to handle increased search spaces is the application of multiple identification steps in general, independently of target application such as strain-level identification. These multi-stage identification search strategies are described by several different terms such as serial search ^23^, multi-step ^24^, iterative ^25,26^, multi-stage ^27^, two-step ^28–30^ as well as cascade search ^31^ and they find application in proteomics, metaproteomics ^26,28^ and proteogenomics ^30,32^. Most of these strategies do not only overlap in their objective of increasing the identification rate or identification confidence, but share methodological principles as well. This includes the concept of identifying primarily unassigned spectra using databases of increasing complexity (for instance, by employing altered digestion parameters, additional post-translational modifications or additional spectral and genomic databases) ^24–27,31^ as well as the recurring theme of database size reduction ^24,26,28–30,32^. In addition, some methods rely on spectral quality assessment to enhance subsequent identification steps ^25,32^ or exhibit a focus on algorithmic runtime reduction ^24^.

Apart from database size, multi-proteome databases present an additional challenge for taxonomic assignment in the form of high sequence similarity between proteomes, in particular between related species and strains. These similarities give rise to taxonomically ambiguous, widespread and false assignments and thus need to be accounted for. Several methods exist that provide various strategies to account for ambiguous peptide spectrum matches due to sequence similarity. Dworzanski et al. apply a proteogenomic mapping approach in combination with discriminant analysis to infer most likely bacterial assignments ^33^. The proteogenomic approach was further developed in BACid that applies several statistical measures to account for similarity, including the comparison of ratios of taxonomic differences and of unique peptides to known error rates, i.e. the noise levels ^34,35^. BACid has since been applied to and extended for samples of unknown origin ^36^ as well as mixtures of bacteria ^36,37^. With the focus on metaproteomic analysis, MEGAN ^38^, UniPept ^39^, MetaProteomeAnalyzer ^40,41^ and TCUP ^6^ rely on lowest common ancestor or most specific taxonomy approaches to assign peptides spectrum matches to taxonomic levels. MiCId uses a clustering approach based on peptidome similarity of taxa on different levels in combination with unified E-values to infer statistically significant representatives of clusters as identified or classified microorganisms ^13,42^. Tracz *et al.* use a direct assignment strategy by concatenating all proteins of a proteome into one pseudo-polyprotein and considering only the top-scored spectrum matches to a peptide as counts for bacterial candidates ^43^. In contrast, Pipasic explicitly makes use of ambiguous peptide spectrum matches (PSMs) by applying abundance correction based on intensity-weighted proteome similarities of organisms in metaproteomic samples ^44^. While these methods share the objective of reducing uncertainty in taxonomic identification due to ambiguous assignments in the context of the protein inference issue ^45^, their application for untargeted strain-level identification is not straightforward for several reasons. For example, unique peptides are not necessarily available for all candidate strains in the sequence databases for which the information given is often limited to species or higher taxonomic levels. In addition, the approaches rely on fully sequenced genomes (and thus uniform proteome coverage) or other forms of targeted, curated, inhouse or validated databases and are thus restricted to a predefined limited set of organisms, often with definite focus on bacteria. Furthermore, some methods do not scale well with immoderate large databases (i.e. high number of proteomes) with respect to computational or analytical performance.

Previous multi-step procedures illustrate the effect of database size on identification confidence and the advantage of applying concise, prefiltered or specialized databases. Thus, we transfer the general concept of multi-step procedures to approach the increased search space necessary for untargeted and detailed taxonomic classification. We present TaxIt, an iterative workflow for untargeted strain-level identification of protein samples. By applying two separate identification steps for species- and strain-level classification, we circumvent the immediate need for a comprehensive database containing strain proteome sequences. Thereby, a first untargeted search allows the selection of a relevant species and enables to focus a second search on a highly reduced but adequate choice of automatically downloaded strain proteomes, resulting in increased identification confidence and reduced taxonomic ambiguity. Moreover, strain-level identification relies on recent data, without the need of keeping a local comprehensive strain database up-to-date. The workflow takes advantage of a free, publicly available and continuously growing protein sequence resource (NCBI Protein) and is thus suitable for most common established MS experimental workflows.

## Material and Methods

The TaxIt workflow for strain identification consists of several recurring modules interconnected, controlled (in terms of input and output) and automatically executed by the Snakemake computational workflow management system ^46^. The designed workflow executes up to three stages including (i) an optional host filter, (ii) species identification (primary iteration) and (iii) strain identification (secondary iteration). A concise overview of the workflow is illustrated in Figure 1. The main steps of each iteration are the execution of a peptide search engine, false discovery rate (FDR) control, taxonomic classification, taxa counting and adjustment as well as candidate selection and visualization. The automated download of strain proteomes bridges the primary and secondary iteration. The procedures of the main modules are described in detail below.

**Figure 1.**
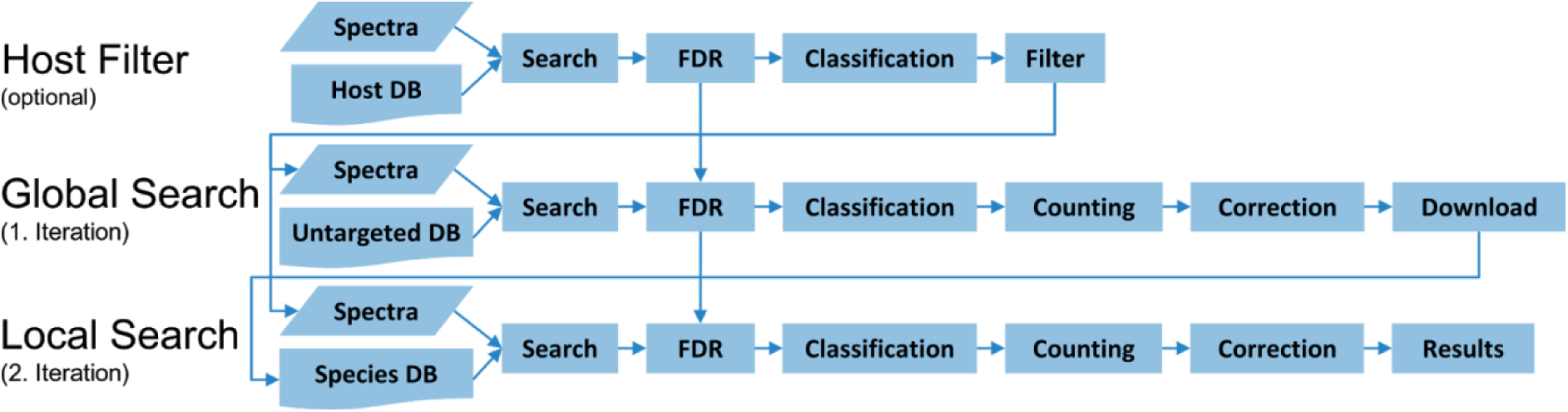
Overview of the TaxIt workflow. In up to three stages, MS/MS spectra are searched against a host proteome to pre-filter host proteins (optional), against an untargeted database for global species identification (primary iteration) and against a targeted, species-based and automatically fetched database for strain level identification (secondary iteration).

### Peptide Search

The central step in the analysis of MS/MS spectra is an efficient peptide search for comprehensive protein identification. Therefore, we rely on established and reliable open-source database search engines such as X!Tandem ^47^ in combination with the XTandem Parser ^48^ and MS-GF+ ^49^. However, any command-line search engine including proprietary ones could be implemented via additional Snakemake rules. We apply a classic target-decoy approach for false discovery rate (FDR) control ^50^. Decoy sequences are created upfront and independently of the search engine with fasta-decoy.pl ^51^ by reversing the target sequences (including contaminant proteins, for instance cRAP ^52^) and both target and decoy databases are concatenated as suggested by Jeong et al. ^20^. PSMs are subjected to an FDR cutoff based on a per-match FDR calculated as N_decoy_/N_target_, with N_decoy_ being the number of decoys in between targets (N_target_) in a list of matches sorted by e-value ^20,53^. To acknowledge established false positive hits of previous iterations, decoy sequences with an e-value above the FDR-cutoff are passed on and concatenated to the database of the next iteration.

### Taxonomic Classification

We make use of the NCBI Taxonomy to assign PSMs to corresponding taxa and redistribute shared hits introduced by proteins associated with higher/upper taxonomic levels such as genus. First, NCBI protein accessions are mapped to NCBI taxids using the NCBI protein id mapping file (ftp.ncbi.nih.gov/pub/taxonomy/accession2taxid/prot.accession2taxid.gz). Next, taxonomic relations are inferred from the NCBI Taxonomy nodes dump file (ftp.ncbi.nih.gov/pub/taxonomy/taxdump.tar.gz) and PSMs are reassigned to a taxonomic level based on the objective of the current iteration. In general, PSMs assigned to higher taxa such as genus are propagated as shared hits to corresponding leaf taxa as long as the target leaves already exhibit matches on their own. In addition, in the first iteration, PSMs are summarized at the species level.

### Count Adjustment

For each candidate taxon (i.e. species in primary and strains in secondary iteration, respectively), raw counts are calculated by summing over all assigned PSMs including non-unique matches. To account for taxonomic biases caused by shared matches, we integrate a weighting scheme based on the level of uniqueness using the global frequency of a PSM. Here, a PSM count per taxon is adjusted by the number of occurrences in all candidate taxa. Thereby, unique PSMs gain in value for the final taxa selection without fully neglecting the importance of high numbers of non-unique matches that are often highly present within closely related taxa such as strains.

### Selection and Downloads

After count adjustment, the most dominant candidate taxon is selected as the most likely species or strain, respectively. In the final step of the first iteration, the selected species is used to retrieve strain-level data for the strain identification in the second iteration. Once more, we rely on the NCBI Taxonomy and infer all available strains for the candidate species via the nodes dump file. Next, strain proteins are automatically downloaded from NCBI Protein using the NCBI Entrez API ^54^ in combination with the jsoup: Java HTML Parser ^55^. This includes all available RefSeq as well as non-RefSeq sequences since the availability of curated strain material is often limited. Finally, the obtained protein sequences are merged into a single database and redundant entries are removed using seqkit rmdup ^56^.

### Benchmarking Experiments

For benchmarking, we compare TaxIt against classic comprehensive search strategies based on straight non-iterative taxonomic identification supported by unique PSMs or abundance similarity correction as provided by Pipasic ^44^.

TaxIt uses NCBI RefSeq proteins of selected kingdoms as reference databases for initial species identification followed by automated and selective strain protein incorporation. Unique-PSMs- and Pipasic-based strategies, however, apply comprehensive databases integrating as many strain-level sequences as possible at once, including all protein sequences from the NCBI Protein database for selected kingdoms. In general, a preselection of kingdoms may be justified by clinical findings based on, for instance, symptoms or microscopic examination ^57^. Both, unique-PSMs- and Pipasic-based strategies use the same procedures for peptide search, FDR control and taxonomic classification as described in the iterative workflow. However, PSMs are not summarized at species level and counts are directly inferred at the lowest possible taxonomic level. For the unique-PSMs-based strategy, adjusted counts are based on PSMs that occur only once. In other words, only spectra assigned unambiguously to only one peptide sequence and thus organism are taken into account. The abundance similarity correction of Pipasic uses the similarity of expressed proteomes between taxa to account for attribution biases. Originally intended for metaproteomic abundance correction, it is here applied to highlight the most likely strain within a provided sample.

Since Pipasic is sensitive to a high number of taxa, we limit the input to taxa with a minimum of two hits as well as to the 100 most abundant taxa. Expressed proteins per taxa are extracted according to the taxonomic classification and digested peptides with a minimum length of six amino acids are prepared using trypsin digestion ^58^. PSMs and tryptic peptides are then passed to Pipasic to obtain corrected relative counts.

We perform all three strategies on several viral and bacterial MS/MS spectra samples with available strain-level knowledge. This includes a *Cowpox virus (Brighton Red)* strain (acquired in-house), an *Avian infectious bronchitis virus (strain Beaudette CK)* ^59^ and a *Bacillus subtilis BSN238* ^60^ that we will refer to as cowpox, bronchitis and bacillus sample, respectively. *B. subtilis BSN238* is a transgenic organism resulting from horizontal gene transfer (HGT) of the DivIVA protein from *Listeria monocytogenes strain EGD-e* to *Bacillus subtilis* subsp*. subtilis str. 168*. Since *B. subtilis BSN238* is neither yet present in the NCBI Taxonomy nor in the NCBI Protein database and only one protein is modified, we expect *B. subtilis* subsp*. subtilis str. 168* to be selected as final strain candidate. Bacillus samples are examined twice, once completely with 28902 spectra (bacillus all) and once randomly reduced to 1000 spectra (bacillus 1k) to improve performance with respect to the vast bacterial search space. A detailed description of the cowpox sample acquisition and the search parameters for all samples is provided in Supplementary item 1.

For the viral samples we used all viral NCBI RefSeq proteins (314156 entries, via Entrez on July 10, 2017) in the iterative workflow and additionally all viral non-RefSeq proteins (1987275 entries in total, via Entrez on July 11, 2017) for the unique-PSMs- and Pipasic-based strategies. Viral spectra were filtered beforehand using corresponding host proteomes (all isoforms) including the UniProt *Homo sapiens* reference proteome (93588 entries, UP000005640, May 23, 2017) for the cowpox sample and *Gallus gallus Red jungle fowl* reference proteome (29732 entries, UP000000539, May 16, 2017) for the bronchitis sample. For the iterative examination of the bacillus samples, all bacterial NCBI RefSeq proteins (65996005 entries, release 82) were downloaded via FTP (ftp.ncbi.nlm.nih.gov/refseq/release/bacteria/). Since downloading all NCBI Protein entries for bacteria (taxid 2) is impractical and requires too much time using the Entrez API, we used the NCBI Blast NR database as most comprehensive, common and readily available protein resource. This database can be obtained in fasta format via FTP (ftp.ncbi.nlm.nih.gov/blast/db/FASTA/, release 19.07.2017) and the bacteria subset (93760438 entries) was extracted with in-house scripts to create the databases for the unique-PSMs- and Pipasic-based strategies.

## Results

To demonstrate the potential of iterative strain-level identification, we compared TaxIt against classic comprehensive search strategies based on non-iterative taxonomic identification supported by either unique PSMs or the abundance similarity correction of Pipasic ^44^. The final selections of the top taxa candidates for all samples and all three compared identification strategies are summarized in Table 1. For the cowpox sample, identification results agree the most. TaxIt (Figure 2) and Pipasic (**Supplementary item 1 - Figure S1**) are both able to identify the expected *Cowpox virus (Brighton Red)* strain. However, unique PSMs are limited to the parent *Cowpox virus* species and not available at the strain level. For this reason, an incorrect identification of *Bat astrovirus Hil GX bszt12* could occur based on a single unique PSM (**Supplementary item 1 - Figure S1**). The results also show that the original TaxIt counts are not an appropriate measure to distinguish the expected strain from competing candidates (Figure 2). However, the weight-based correction method implemented in TaxIt resolves the present tie and emphasizes the correct *Cowpox virus* strain (*Brighton Red)*.

**Table 1.**
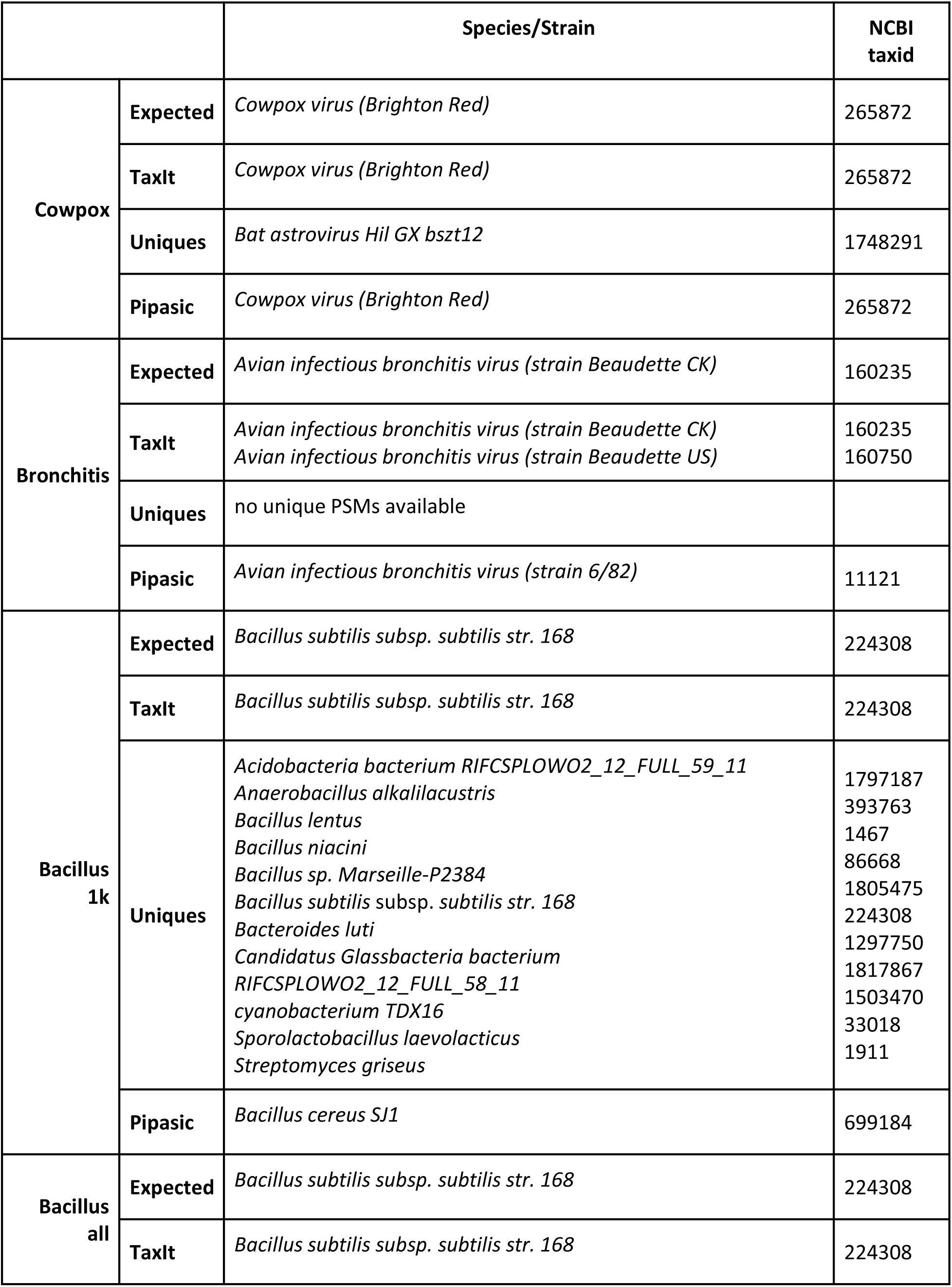
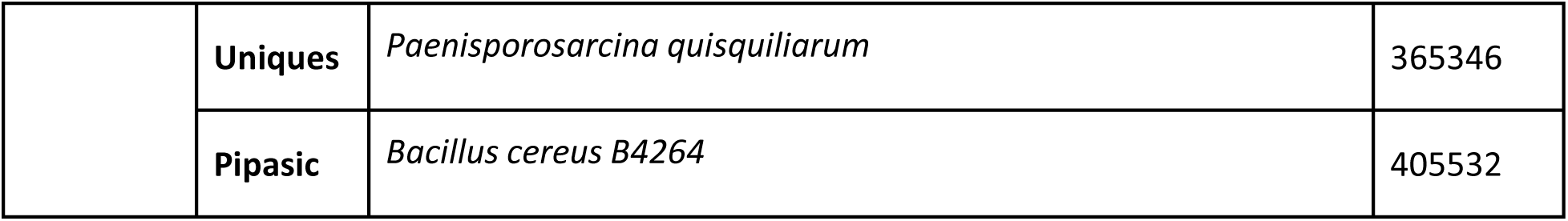
Expected and identified taxa per sample and strategy.

**Figure 2.**
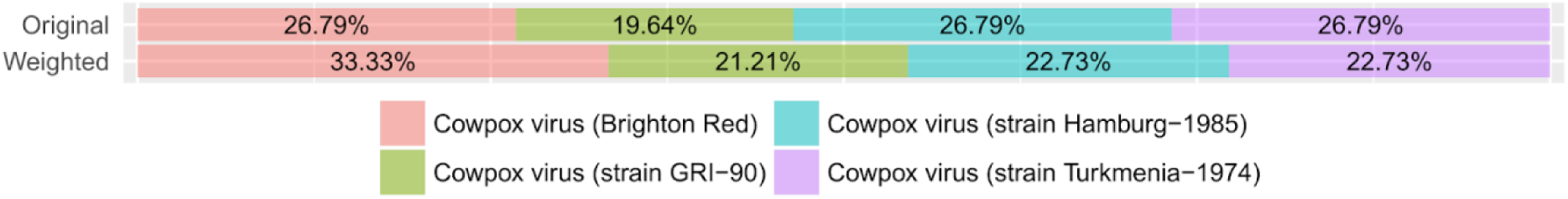
Relative counts of cowpox sample analysis. Relative counts are illustrated as result of the TaxIt workflow. Original and weighted relative counts are summarized by means of one vertical stacked bar each. Candidate strains are labeled and color-coded and ratios highlighted as percentages within bars.

In comparison, bronchitis and bacillus samples feature notable variability in proposed taxa candidates. For bronchitis, TaxIt could identify the expected *Avian infectious bronchitis virus (strain Beaudette CK)* strain. Despite applying a small correction based on weighting, however, final candidate selection is not limited to a single strain but additionally includes a closely related strain, namely *Avian infectious bronchitis virus (strain Beaudette US)* (Figure 3A). Nevertheless, TaxIt provides a highly constrained selection of candidates in the first place. In contrast, the Pipasic-based strategy results in considerably more initial candidates and eventually promotes an incorrect strain, namely *Avian infectious bronchitis virus (strain 6/82)* (Figure 3B). Since no unique PSMs are available for the bronchitis sample, strain-level identification was not possible with this strategy.

**Figure 3.**
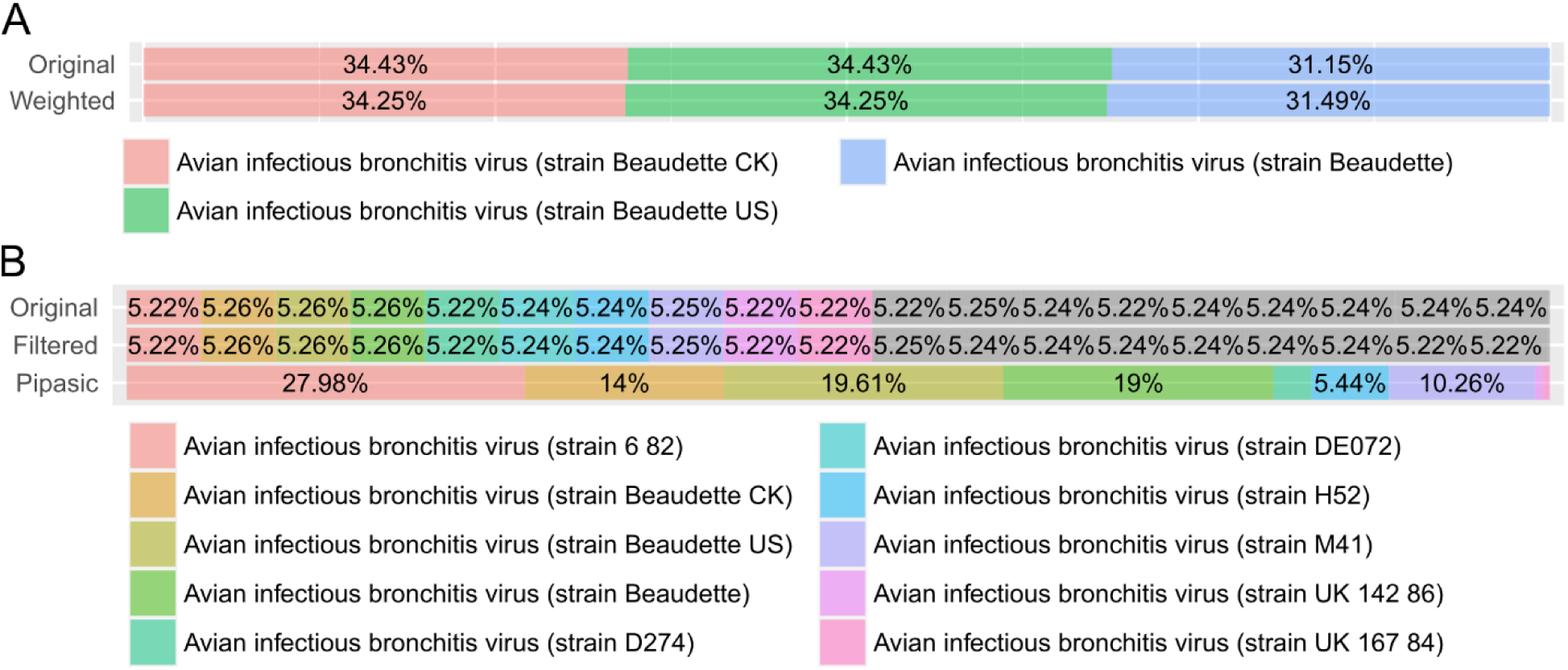
Relative counts of bronchitis sample analysis. Relative counts are illustrated as result of the TaxIt (A) or Pipasic approach (B), respectively. TaxIt’s original and weighted relative counts as well as Pipasic’s original, filtered (a minimum of two hits and only the most 100 abundant taxa) and corrected relative counts are summarized by means of one vertical stacked bar each. Candidate strains are labeled and color-coded and ratios highlighted as percentages within bars.

**Figure 4.**
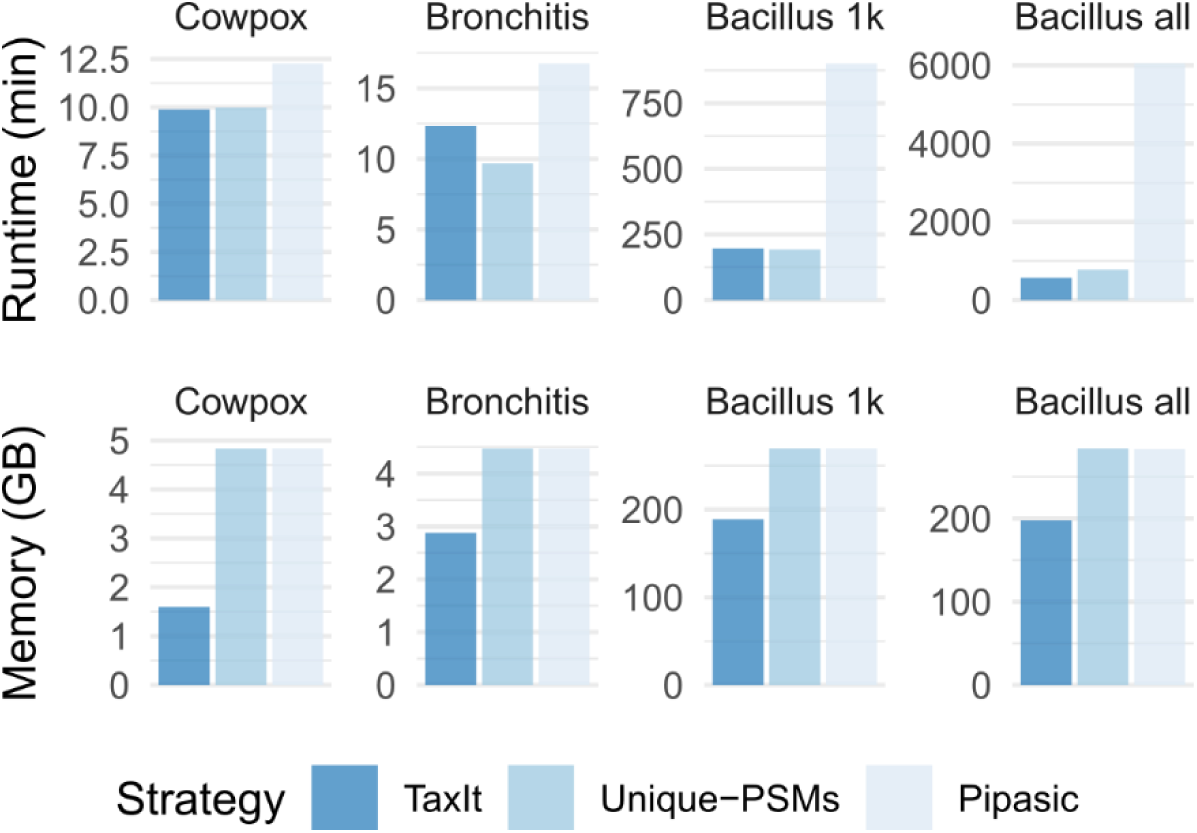
Runtime and memory benchmarks. Total runtime in minutes and maximum memory usage in gigabyte is represented per sample and color-coded identification strategy.

Using TaxIt, the expected *B. subtilis str. 168* strain is identified correctly in the reduced as well as in the complete sample (**Supplementary item 1 - Figure S3 and S4**). Pipasic consistently misses the correct strain and species in favor of incorrect strains such as *Bacillus cereus SJ1* and *Bacillus cereus B4264*. The unique-PSMs-based strategy includes the correct strain *B. subtilis str. 168* into the final candidate list of the bacillus 1k sample. However, it fails to separate the strain from several distinct species due to equal amounts of unique PSMs (**Supplementary item 1 - Figure S3**). Furthermore, the data analysis on the complete bacillus sample results in the incorrect species *Paenisporosarcina quisquiliarum* being predominantly present in terms of uniquely assigned PSMs.

Besides identification efficiency, computational performance may delimit the potential application as routine or expeditious method in research or clinics. Therefore, we compared runtime and memory consumption for all samples and strategies (as illustrated in Table 2 and **Figure 5**). Analysis was performed with X!Tandem as database search engine and limited to 24 threads on a server with Debian GNU/Linux 8.9 (jessie), 64 cores (128 threads) of type Intel(R) Xeon(R) CPU E5-4667 v4 @ 2.20 GHz, 512 GB of RAM and SSD storage. Applying the iterative approach reduces memory usage down to one third for viral strain identification and two third for bacterial strain identification. While the runtime of Pipasic is comparably high, TaxIt shows no substantial change in runtime in comparison to the unique-PSMs-based strategy for small databases such as the collective viral sequences or small sample sizes. However, analyzing the full bacillus sample reveals an increased runtime when using NCBI RefSeq proteins plus selected strain proteomes instead of the extensive NCBI Blast NR database.

**Table 2.** Runtime and maximal memory consumption (resident set size, RSS) per identification strategy and sample.

|  | Runtime (h:m:s) |  |  | Memory (max RSS MB) |  |  |
| --- | --- | --- | --- | --- | --- | --- |
|  | TaxIt | Uniques | Pipasic | TaxIt | Uniques | Pipasic |
| **Cowpox** | 0:09:53 | 0:09:59 | 0:12:16 | 1638.28 | 4949.69 | 4951.09 |
| **Bronchitis** | 0:12:20 | 0:09:41 | 0:16:45 | 2946.19 | 4579.61 | 4579.84 |
| **Bacillus 1k** | 3:15:59 | 3:12:41 | 15:01:04 | 193357.94 | 276134.38 | 276134.28 |
| **Bacillus all** | 9:29:06 | 13:00:55 | 125:53:07 | 202574.46 | 290719.44 | 290487.57 |

## Discussion

TaxIt demonstrates superior performance with respect to strain-level identification and computational expense. In summary, our approach can be used to unambiguously identify candidate organisms of all samples down to a low taxonomic level, with the minor exception of a tie for the bronchitis sample. In contrast, the unique-PSMs-based identification strategy is repeatedly deficient at the strain level or features highly ambivalent results. Furthermore, Pipasic frequently favors an incorrect strain or even an incorrect species. For some samples, correct strains are observed as a top candidate even in the original counts, independently of database choice and prior to count adjustments. Nevertheless, all samples benefit either from the iterative and focused database usage, the count adjustment procedure or the reduced resource consumption while the outcome remains legitimate.

It was shown that TaxIt improves taxonomic assignment already in the count-based ranking, independently of count adjustments, by limiting candidates to strains of a single species early on (Figure 3). For instance, the search of bronchitis samples against the NCBI Blast NR bacteria subspace results in a rather uniform distribution of original counts for *Infectious bronchitis virus* strains: the strains Beaudette, Beaudette CK and Beaudette UK are only slightly increased in comparison to other strains (Figure 3B, **Original**). In contrast, the iterative approach results in less strain candidates of the same species focusing solely on Beaudette strains when selectively searching against *Avian coronavirus* strains in the secondary iteration (Figure 3A, **Original**). This is a consequence of how and whether parental or multispecies proteins are associated with strains or taxa in general within the NCBI taxonomy. In case of the iterative approach, fewer mutual species or genus proteins are consulted in the strain identification iteration. However, the extent of this effect varies between samples, candidate strains or target databases and taxonomies, respectively. For instance, bacterial strain proteomes such as the *Bacillus subtilis* strains feature numerous directly assigned mutual protein sequences and thus result in an extended range of candidate strains of the same species (**Supplementary item 1 - Figure S3 and S4**).

Nevertheless, the restriction to a specific set of strain proteomes prevents manifold primary mis-assignments to distant species, genera or even phyla as can be observed for Pipasic- and unique-PSMs-based original counts, respectively, that cannot be sufficiently resolved after correction (**Supplementary item 1 - Figure S3 and S4**). In general, the iterative and selective database usage ensures that final strain selection is limited to strain candidates of an appropriate species. Thus, it prevents false positive hits on distinct strains of other taxa including species, genera and phyla and allows for a more confident final strain candidate selection.

Furthermore, uniform distributions and even consensus in original counts of strain candidates demonstrate the need and benefit of count adjustment methods. The implemented weighting procedure can resolve ties between strains such as in the TaxIt cowpox sample analysis (Figure 2) or at least amplifies the correct strain and increases the distance to competing candidates.

TaxIt infers exactly one strain for the presented samples each expect for the bronchitis sample where the strains Beaudette CK and Beaudette UK cannot be differentiated. We observed that the corresponding PSMs are fully shared between the two proteomes. Although different proteins are available for each strain in general, peptide hits are either assigned to shared proteins of the parent strain *Infectious bronchitis virus* or to homologous proteins that differ only in identifier but not in sequence. Though a more granular taxonomic relation cannot be ascertained from the NCBI taxonomy, we expect the Beaudette strains to feature a considerably closer relationship as compared to other *Infectious bronchitis virus* strains. Therefore, we consider the draw between Beaudette strains as sufficiently appropriate strain identification.

For the unique-PSMs-based strategy, we observed a poor availability of unique PSMs at the strain level. While the exploitation of purely unique features is a common theme for species-level identification, the low amount of unique PSMs in strains is insufficient for strain-level inference. However, the frequency of spectra matching to distinct proteins and proteomes remains a valuable parameter for strain differentiation when considering and weighing both, unique and the plethora of non-unique matches.

TaxIt has a comparable computational runtime for small samples and databases (such as the viral data) despite using a constrained search space. This is primarily a result of the additional strain proteome downloads, since the NCBI Entrez API is not designed and optimized for large scale downloads and proteins need to be fetched in numerous iterations of small chunks. However, the download overhead fades into the background when considering full bacterial samples such as bacillus all (Table 2 and **Figure 5**) and gives place to a runtime improvement of three quarters when compared to NCBI Blast NR database searches. In contrast, the runtime of Pipasic is afflicted with additional sequence comparisons necessary for constructing the similarity matrix that is highly influenced by increasing numbers of PSMs and taxa to compare. Finally, the memory footprint of TaxIt in comparison to the unique-PSMs- and Pipasic-based strategies remains constantly less for all samples, as would be expected when using substantially less proteins in the search databases.

On a final note, strain-level identification performance is generally limited by the availability and integrity of taxa and proteomes in used databases. However, constantly increasing quality and quantity of the NCBI Taxonomy and Protein databases will induce constant improvement of strain-level identification strategies such as the presented iterative workflow.

## Conclusion

Untargeted strain level identification via MS/MS spectra is a challenging task with respect to the excessive quantity of strains that need to be considered competitively. For solving this problem, we present TaxIt, an iterative approach that first focuses on species identification and thus limits strain identification to concise selected target databases. In our method, both iteration steps take advantage of publicly available data from the NCBI Taxonomy and Protein databases. Thereby, TaxIt enables final strain identification from recent and relevant sequence data, without the need of heavily curated, tailored or taxonomically constrained databases. TaxIt supports any MS/MS peptide data suitable for classic database search methods. With further ongoing developments, our tool might enable a wide range of applications in public health, research and potentially also clinics in the future: based on further validation, TaxIt might support diagnosis and treatment of pathogen-based infectious diseases. TaxIt is available for download under open-source license at https://gitlab.com/rki_bioinformatics.

## Supporting information

Supplementary item 1

## Author Contributions

MK carried out the implementation and evaluation, participated in the software design and experimental design and drafted the manuscript. JD participated in the experimental design and data acquisition. AN participated in the experimental design. TM participated in the software design and drafted the manuscript. BYR participated in the software design and experimental design and drafted the manuscript. All authors read and approved the final manuscript.

## ACKNOWLEDGMENT

The authors would like to thank Henning Schiebenhoefer for constructive criticism of the manuscript.

This work was supported by the German Research Foundation (DFG) (RE3474/2-2 to B.Y.R.).

## ABBREVIATIONS

FDR: false discovery rate
LC: liquid chromatography
MALDI: matrix-assisted laser desorption/ionization
MS: mass spectrometry
PSM: peptide spectrum match
RSS: resident set size
TOF: time-of-flight

