## Supplementary item 1 for "An iterative and automated computational pipeline for untargeted strain-level identification using MS/MS spectra from pathogenic samples"

### S1 Search Parameters

#### Cowpox Sample

The cowpox sample of strain *Cowpox virus (Brighton Red)* was acquired in-house, as described in *S2 Cowpox Sample Acquisition*. The mass spectrometry proteomics data have been deposited to the ProteomeXchange Consortium via the PRIDE [1] partner repository with the dataset identifier PXD014913. Spectra were analyzed applying a tryptic search with parent ion mass tolerance of 10 ppm, fixed modification cysteine carbamidomethylation (+57 Da) as well as an additional variable modification methionine oxidation (+16 Da).

#### Bronchitis Sample

Bronchitis samples of the strain *Avian infectious bronchitis virus (strain Beaudette CK)* were downloaded from PRIDE (PXD002936) and the sample "BeauR2.raw" was randomly selected for analysis. The raw file was converted to an mgf file using ProteoWizard's MSConvert GUI (3.0.8764) [2]. Spectra were analyzed with default settings including a tryptic search with fixed modification cysteine carbamidomethylation (+57 Da) and parent ion mass tolerance of 100 ppm.

#### Bacillus Sample

The bacillus sample of the strain *Bacillus subtilis* subsp. *subtilis str. 168* was download from PRIDE (PXD007242, file "614_NG4_BSN238_Urea-Trp_1ug_SR-LFQ_4h_161201.mgf").
Spectra were analyzed with default settings including a tryptic search with fixed modification cysteine carbamidomethylation (+57 Da). However, parent ion mass tolerance was set to 10 ppm in accordance with the original publication.

[1] Vizcaíno, J. A.; Csordas, A.; del-Toro, N.; Dianes, J. A.; Griss, J.; Lavidas, I.; Mayer, G.; Perez-Riverol, Y.; Reisinger, F.; Ternent, T.; et al. 2016 Update of the PRIDE Database and Its Related Tools. Nucleic Acids Res 2016, 44 (D1), D447–D456. https://doi.org/10.1093/nar/gkv1145.
[2] Chambers, M. C.; Maclean, B.; Burke, R.; Amodei, D.; Ruderman, D. L.; Neumann, S.; Gatto, L.; Fischer, B.; Pratt, B.; Egertson, J.; et al. A Cross-Platform Toolkit for Mass Spectrometry and Proteomics. Nat Biotech 2012, 30 (10), 918–920. https://doi.org/10.1038/nbt.2377.

### S2 Cowpox Sample Acquisition

#### CPXV-infection

One day prior infection, 5x105 HEp-2 cells were seeded into 6 well plates with 4 mL cell culture medium (DMEM supplemented with 10 % FCS and 2 mM L–Glutamine) each and kept in an incubator at 37°C for 16 h. The medium was removed and cells were infected with CPXV Brighton Red (BR) (ATCC® VR-302™) using a multiplicity of infection (MOI) of 0.1 in 1 mL cell culture medium per well for 1 h. Again, the medium was removed and cells were washed once with 5 mL phosphate-buffered saline (PBS) before they were further incubated in 4 mL cell culture medium for 24h. Supernatant was removed and mixed 1:1 with 4 % SDS, 0.1 M Tris-HCl (pH 7.6), 10 mM Tris(2-carboxyethyl)phosphine (TCEP) and 40 mM 2-chloroacetamide (CAA) at 99°C for 5 min. DNA was sheared by sonication for 3 x 1 min on ice using a 2" Sonifier™ Cup Horn in a Cell Disruptor (Branson Ultrasonics Corporation, Danbury, CT, USA) and lysates were clarified by centrifugation at 16,000 × g for 10 min.

#### Filter-aided sample preparation (FASP)

Samples were prepared using the filter-aided sample preparation (FASP) protocol with minor modifications. Briefly, 20 µg of each sample were processed using Vivacon 500 Centrifugal Ultra Filter (Sartorius, Goettingen, Germany) with a molecular weight cut-off (MWCO) of 30 kDa. SDS was depleted 4 x with 200 µL 8 M Urea in 50 mM Tris-HCl, pH 8.5, before the urea concentration was reduced using 3 x 100 µL 50 mM Tris-HCl, pH 8.5. The proteins were digested for 16 h at 37°C in 50 mM Tris-HCl, pH 8.5 using Trypsin Gold, Mass Spectrometry Grade (Promega, Fitchburg, WI, USA) at a protein/enzyme ratio of 100:1. The peptides were collected by centrifugation and two washing steps with 40 µL 50 mM Tris-HCl, pH 8.5. The peptides were desalted using 200 µL StageTips packed with four Empore™ SPE Disks C18 (3M Purification, Inc., Lexington, USA) and concentrated using a vacuum concentrator but not dried completely. Samples were filled up to 16 µL with 0.1% formic acid and peptides were quantified by measuring the absorbance at 280 nm using a Nanodrop 1000 (Thermo Fisher Scientific, Rockford, IL, USA).

#### nLC-MS/MS

Single-run bottom-up proteome analysis was performed on an Easy-nanoLC (Proxeon, Odense, Denmark) coupled online to an LTQ Orbitrap Discovery™ mass spectrometer (Thermo Fisher Scientific, Rockford, IL, USA). 2 µg peptides were loaded directly on a Reprosil-Pur 120 C18-AQ, 2.4 µm, 300 mm x 75 µm fused silica capillary column (Dr. Maisch, Ammerbuch-Entringen, Germany), which was kept at 60°C using a butterfly heater (Phoenix S&T, Chester, PA, USA). Peptides were separated using a linear 240 min gradient of acetonitrile in 0.1 % formic acid and 3 % DMSO from 0 to 29 % at 200nL/min flow rate. The mass spectrometer was operated in a data-dependent manner in the m/z range of 400 – 1,400 with a resolution of 30,000 in the orbitrap. Up to the seven most intense 2+ and 3+ charged ions were selected for low-energy CID type fragmentation in the ion trap with a normalized collision energy of 35 % using an activation time of 10 ms. The m/z isolation width for MS/MS fragmentation was set to 2 Th. Once fragmented, up to 500 isolated peaks were dynamically excluded from precursor selection for 90 s within a 20 ppm window. The ion selection threshold for $MS^2$ spectra was 1,000 counts, and the maximum allowed ion accumulation times were 500 ms for full scans and 100 ms for $MS^2$ spectra. Automatic gain control was set to a target value of 1e6 for full scans and 5e3 for $MS^2$. Peptides were ionized using electrospray with a stainless steel emitter, I.D. 30 µm, (Proxeon, Odense, Denmark) at a spray voltage of 2.1 kV and a heated capillary temperature of 275°C. The background ion signal intensities were reduced using an ABIRD device (ESI Source Solutions, Woburn, MA, USA).

### S3 Additional Figures

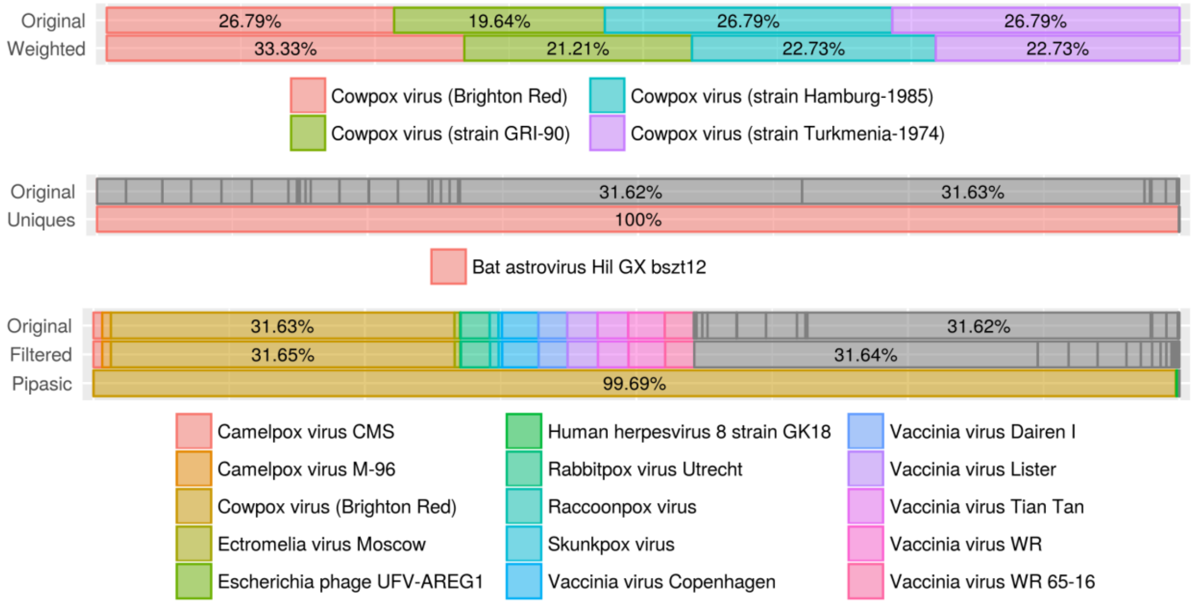


**Figure S1: Relative counts of cowpox.** Relative counts are illustrated for TaxIt (top), uniques- (middle) and Pipasic-based search strategies (bottom). Original, filtered (if applicable) and corrected relative counts are summarized by means of one vertical stacked bar each. Taxa are labeled and color-coded based on a limit of 15 final top candidates (i.e. after correction) with a relative count greater zero. Furthermore, ratios greater than 0.05 are highlighted as percentages within bars. Selecting unique PSMs resulted in only one PSM for *Bat astrovirus Hil GX bszt12* (taxid 1748291).

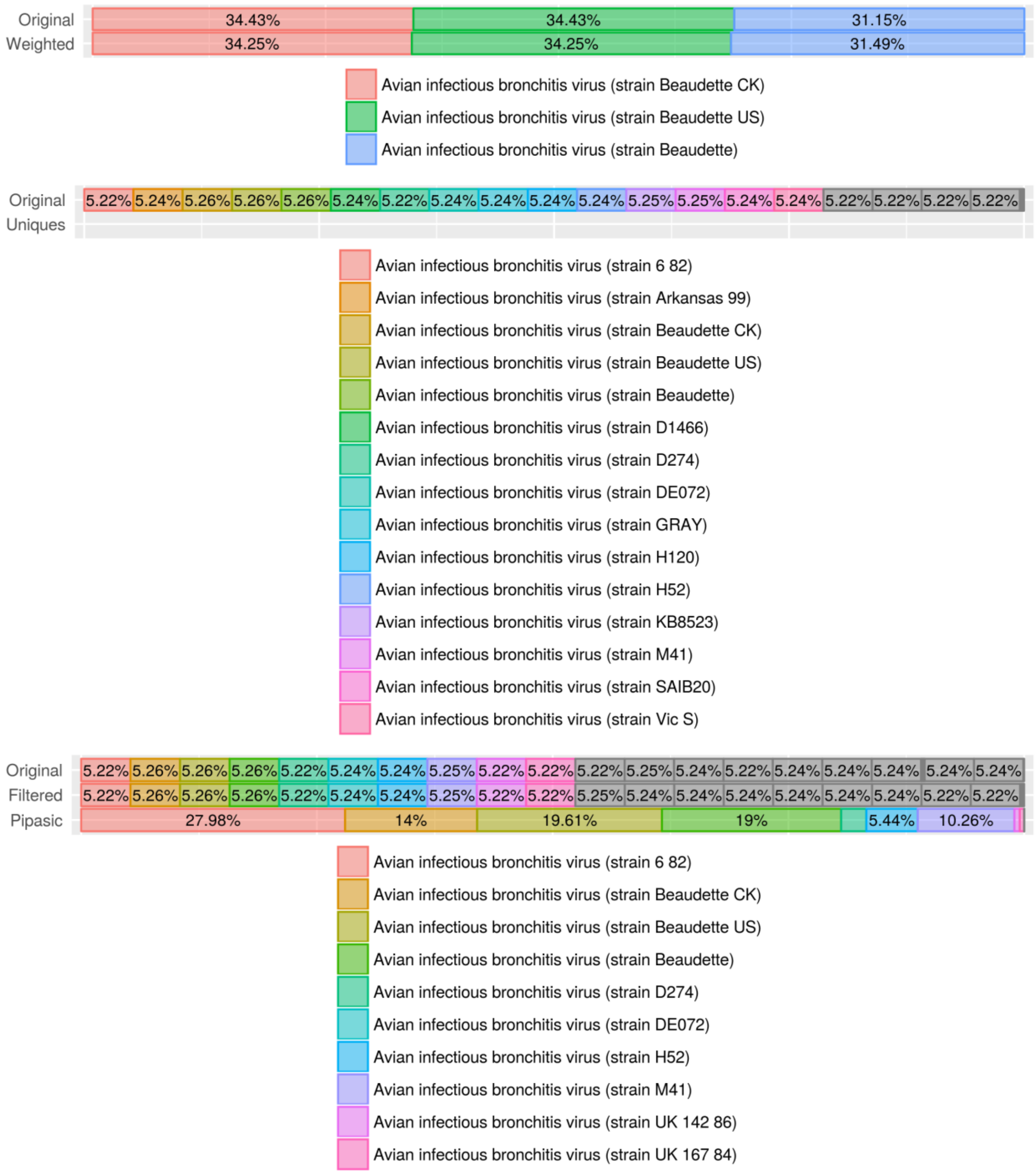


**Figure S2: Relative counts of bronchitis.** Relative counts are illustrated for TaxIt (top), uniques- (middle) and Pipasic-based search strategies (bottom). Original, filtered (if applicable) and corrected relative counts are summarized by means of one vertical stacked bar each. Taxa are labeled and color-coded based on a limit of 15 final top candidates (i.e. after correction, except for uniques) with a relative count greater zero. Furthermore, ratios greater than 0.05 are highlighted as percentages within bars.

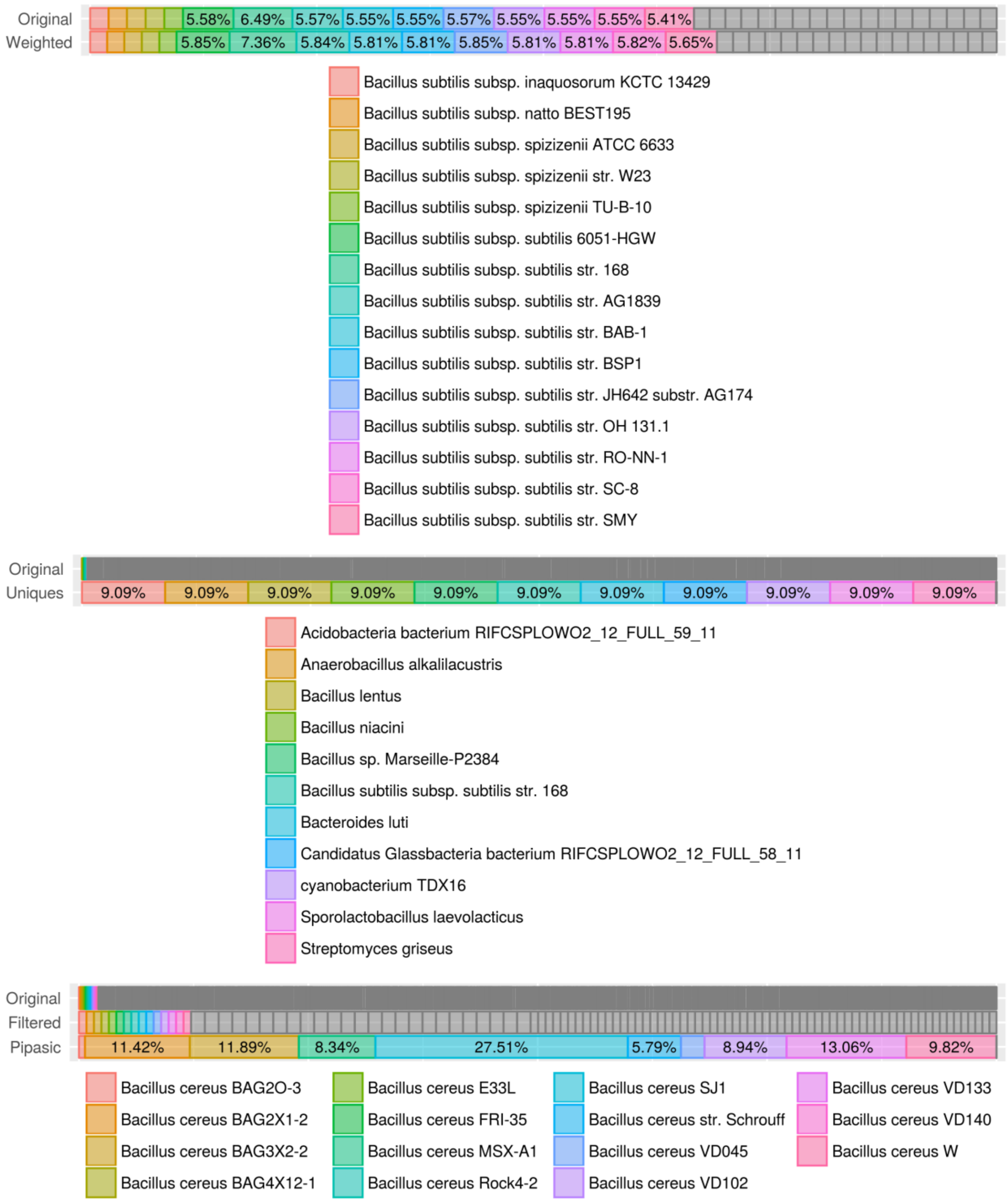


**Figure S3: Relative counts of bacillus 1k.** Relative counts are illustrated for TaxIt (top), uniques- (middle) and Pipasic-based search strategies (bottom). Original, filtered (if applicable) and corrected relative counts are summarized by means of one vertical stacked bar each. Taxa are labeled and color-coded based on a limit of 15 final top candidates (i.e. after correction) with a relative count greater zero. Furthermore, ratios greater than 0.05 are highlighted as percentages within bars.

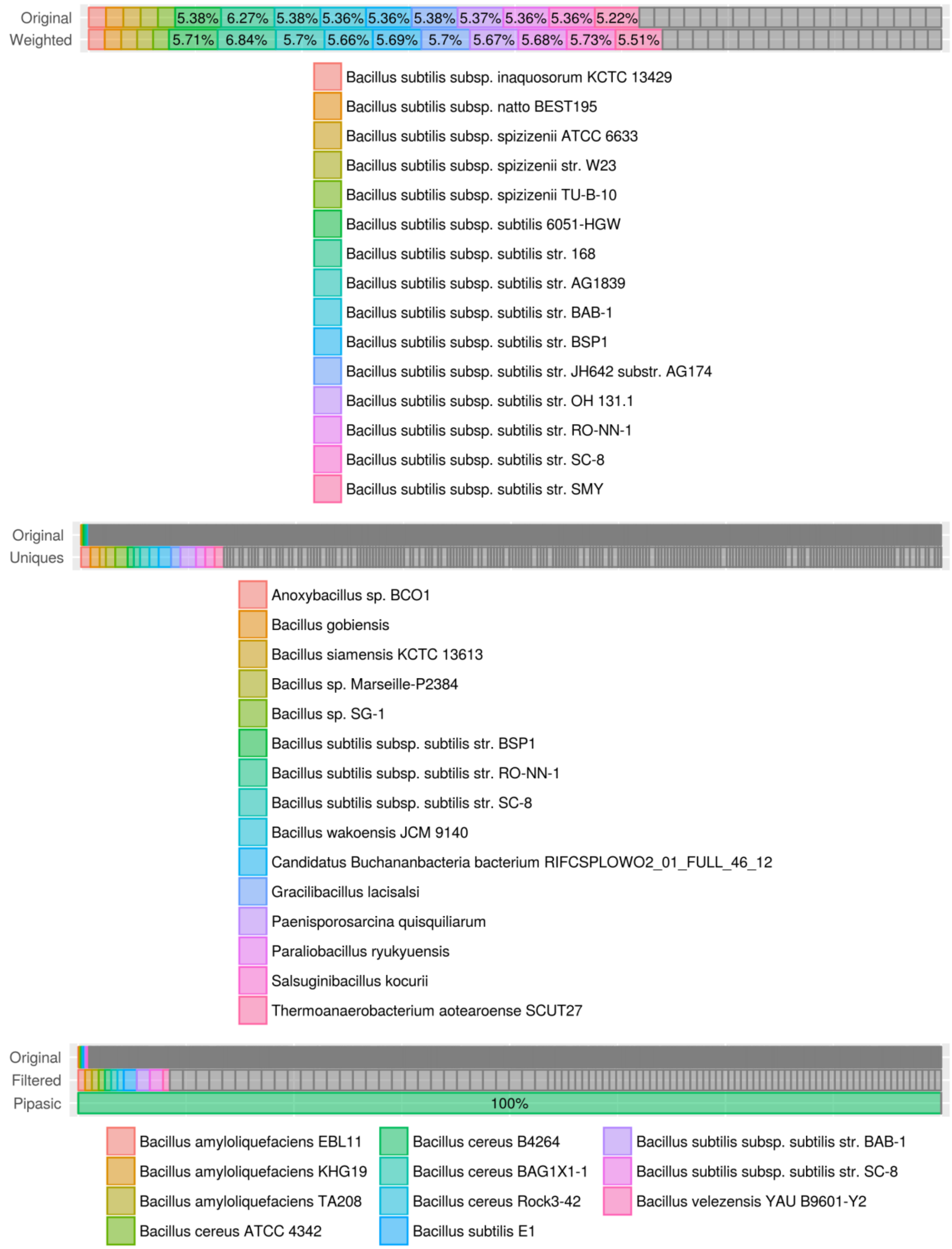


**Figure S4: Relative counts of bacillus all.** Relative counts are illustrated for TaxIt (top), uniques- (middle) and Pipasic-based search strategies (bottom). Original, filtered (if applicable) and corrected relative counts are summarized by means of one vertical stacked bar each. Taxa are labeled and color-coded based on a limit of 15 final top candidates (i.e. after correction) with a relative count greater zero. Furthermore, ratios greater than 0.05 are highlighted as percentages within bars.
